# Defining subpopulations of differential drug response to reveal novel target populations

**DOI:** 10.1101/435370

**Authors:** Nirmal Keshava, Tzen S. Toh, Haobin Yuan, Bingxun Yang, Michael P. Menden, Dennis Wang

## Abstract

Personalised medicine has predominantly focused on genetically-altered cancer genes that stratify drug responses, but there is a need to objectively evaluate differential pharmacology patterns at a subpopulation level. Here, we introduce an approach based on unsupervised machine learning to compare the pharmacological response relationships between 327 pairs of cancer therapies. This approach integrated multiple measures of response to identify subpopulations that react differently to inhibitors of the same or different targets to understand mechanisms of resistance and pathway cross-talk. MEK, BRAF, and PI3K inhibitors were shown to be effective as combination therapies for particular *BRAF* mutant subpopulations. A systematic analysis of preclinical data for a failed phase III trial of selumetinib combined with docetaxel in lung cancer suggests potential indications in urogenital and colorectal cancers with *KRAS* mutation. This data-informed study exemplifies a method for stratified medicine to identify novel cancer subpopulations, their genetic biomarkers, and effective drug combinations.

## Introduction

Drug developers face a conundrum in predicting the efficacy of their investigational compound compared to existing drugs used as the standard of care treatment. Systematic screening of drug compounds across a variety of genomic backgrounds in cancer cell lines has improved clinical trial design and personalized treatments ^1^. Following the pioneering NCI-60 screen comprised of 59 unique cell lines ^2^, modern high-throughput screens such as the Genomics of Drug Sensitivity in Cancer (GDSC) ^3,4^, the Cancer Cell Line Encyclopedia (CCLE) ^5^ and the Cancer Therapeutics Response Portal (CTRP) ^6–8^ have characterised >1,000 cancer cell lines with the goal of establishing the genetic landscape of cancer. The deep molecular characterisation of these large cell line panels is complemented with high-throughput drug screens, which enables the discovery of drug response biomarkers. For example, analysis of the generic BRAF inhibitors PLX4720, SB590885 and CI-1040 reproduced drug sensitivity association with *BRAF* mutation in melanoma, or afatinib sensitivity with *ERBB2* amplifications in breast cancer ^3,4,9^. These associations between genetic variants and treatment response have helped identify specific patient subpopulations who are most likely to benefit from treatment. In Phase III clinical trials, however, for new drugs to be successful, they must demonstrate a significant improvement over the existing standard of care. Accurately defining in which subpopulations a new drug demonstrates improved differential efficacy over other drugs targeting the same disease could lead to both better clinical outcomes as well as new targeted therapies.

While several methods have been proposed to identify drug response biomarkers in cell lines for precision medicine and drug repositioning ^4,5,10,11^, there is a need for more objective and unsupervised approaches for identifying subpopulations with differences in drug response (differential drug response), and consequently systematically gain mechanistic insights from biomarkers. Most approaches capable of comparing multiple drugs measure the overall similarity (or correlation) based on a single response summary metric ^7,12^, which permits drug repositioning based on subpopulations with similar behavior, but neglects ones that behave differently (**Figure S1A**). Here, we used an unsupervised technique to identify the perimeters of differentially sensitive or resistant subpopulations and which may be generalized to stratify the pharmacology response for any pair (or n-tuple) of targets using any number of drug response summary metrics (e.g. IC_50_ or AUC). Segmentation of the overall population occurs top-down and along globally-optimal contours that are derived explicitly and maximize the differences between the two resulting subpopulations. The segmentation continues recursively and is modulated by multiple user-defined criteria such as the size or separability of the resulting subpopulations. Higher threshold values for both result in less granular subpopulations but increase certainty that the subpopulations and the quantities estimated from them are both distinct and accurate.

We present results from our platform, SEABED (SEgmentation And Biomarker Enrichment of Differential treatment response), to demonstrate how unsupervised machine learning can discover intrinsic partitions in the drug response measurements of two or more drugs that directly correspond to distinct pharmacological patterns of response with therapeutic biomarkers. Addressing the challenges in comparing the response of two drugs, SEABED initially assesses two gold standards with established clinical biomarkers, namely the differential response of a BRAF inhibitor and MEK inhibitor with anticipated *BRAF* and *KRAS* mutations ^13–16^, and an EGFR inhibitor and MEK inhibitor with expected biomarkers of *EGFR, ERBB2* and *KRAS* mutations ^17–20^. Next, we systematically compare how different drugs targeting the MAPK and PI3K-AKT pathway yield different patterns of response within subpopulations. We show how differential drug response may indicate benefit for drug combinations explained through independent action rather than probable synergy by examining subpopulations uniquely sensitive to a single drug ^21^, which may be precisely targeted by identified biomarkers. Finally, we demonstrate how the analysis of differential response can guide the design of clinical trials by revealing specific indications where an investigational therapy may be more effective than the standard treatment.

## Results

We applied our technique to discover subpopulations of cell lines in which two or more compounds, possibly addressing the same disease state or even targeting the same genetic alteration, have a common pharmacological pattern of response. By further associating enriched genetic alterations in subpopulations with specific patterns of response, we shed light into molecular mechanisms responsible for patient subpopulations that respond differently to two drugs.

### Identifying subpopulations of differential drug response

We first considered the specific circumstance in which two drugs engage different targets within the same signalling pathway, namely agents targeting MAPK signaling. SEABED used nearly 1,000 cancer cells derived from the GDSC database, and we evaluated two established drug response measures: the drug concentration required to reduce cell viability by half (IC_50_) and the area under the dose-response curve (AUC; **Figure 1A**). SEABED employed a multivariate similarity measure to compare the vector patterns of response for each distinct pair of cell lines without requiring *a priori* assumptions on the number or distribution of the subpopulations. The result is a diverse cell line population segmented into distinct subpopulations having homogeneous patterns of drug response (**Figure 1B**). Here exemplified, we show that the drug response of 802 cell lines treated with either SB590885 (BRAF inhibitor) or CI-1040 (MEK inhibitor) could be segmented into 7 distinct subpopulations with a median size of 40 cell lines by integrating the two metrics of drug response, AUC and IC_50_ (**Figure 1C;** see **Figures S1B and S1C** for individual cell lines segmented by IC_50_ and AUC respectively). We comprehensively evaluate pan-cancer somatic events to nominate biomarkers (**see Methods**) ^4^, and found that the subpopulation sensitive to both inhibitors was significantly enriched for *BRAF* mutants (P=3.87e-14, hypergeometric test), while another subpopulation was exclusively sensitive to the MEK inhibitor and significantly enriched for *KRAS* mutations (P=0.00589, hypergeometric test).

**Figure 1:**
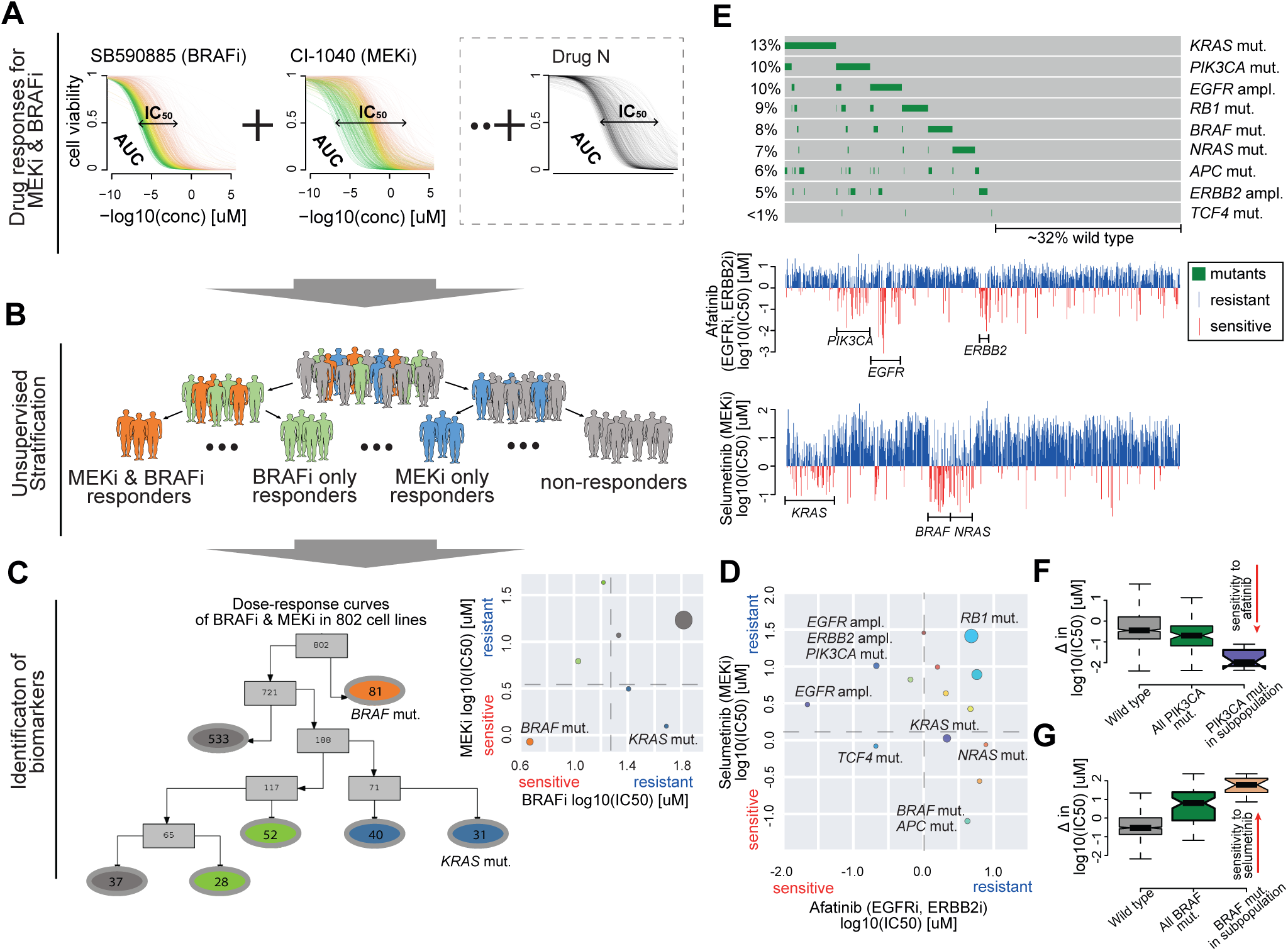
Segmentation of a population based on pharmacological patterns of response discovers subpopulations with differential response. **(A)** Dose response curves of two or more drugs are measured across a population of up to 1,000 cancer cell lines. **(B)** The population is segmented into distinct and homogeneous subpopulations based on their response to multiple drugs. When comparing two drugs, subpopulations can be categorised based on their mean log(IC_50_s): sensitive to both drugs (orange), sensitive to drug A but not drug B (green), sensitive to drug B but not drug A (blue), resistant to both drugs (grey). **(C)** Segmentation results for a BRAF inhibitor (SB590885) and a MEK inhibitor (CI-1040). Tree nodes contain the number of cell lines and are colored based upon their category of response. Significance testing of 735 cancer functional events reveals subpopulations enriched for *BRAF* and *KRAS* mutations. **(D)** Scatter plot showing derived subpopulations based on their pharmacological responses for afatinib and selumetinib. Dashed lines indicate 20th percentile of log(IC_50_) values for each drug. *PIK3CA, EGFR, ERBB2, KRAS, NRAS, BRAF, APC, TCF4* and *RB1* mutations were found enriched in the associated subpopulations. **(E)** OncoPrint visualizing the percentage of mutations of selected genes in cell line panel treated with either afatinib (EGFR inhibitor) or selumetinib (MEK inhibitor). The waterfall plots comparing the response of the cell lines to afatinib and to selumetinib. **(F)** Boxplot of difference in log(IC_50_) values between afatinib and selumetinib response for wild-type cell lines, all cell lines with *PI3KCA* mutation and cell lines in derived subpopulations with enriched *PI3KCA* mutation. **(G)** Same as panel **(F)**, but for *BRAF* mutations.

In another example we examined a case where one inhibitor might overcome resistance to another inhibitor targeting the same pathway; AZD6244/ARRY-142886 selumetinib (MEK inhibitor) with afatinib (EGFR and ERBB2 dual inhibitor) across 839 cell lines (**Figure 1D**). Strong markers of sensitivity for selumetinib are subpopulations carrying known associated *KRAS, NRAS* and *BRAF* mutations (**Figures 1D and 1E**). A less anticipated association is *APC* loss-of-function sensitivity to selumetinib, albeit this was also found with trametinib (another MEK inhibitor) in APC deficient mice ^22^. We reproduced the well-established associations of afatinib with either *EGFR* and *ERBB2* amplifications ^4,23^, and surprisingly our unsupervised segmentation returned two subpopulations enriched for *EGFR* amplifications. The more sensitive subpopulation is solely enriched for *EGFR* amplifications, whilst the less sensitive subpopulation additionally includes activating *PIK3CA* mutations. In concordance with recent literature, PI3K-AKT signaling drives acquired drug resistance to EGFR inhibitors in lung cancer ^24^.

Drug response segmentation resulted in 14 subpopulations with a median size of 37 (**Figure 1D**). The subpopulation enriched for *EGFR, ERBB2* and *PI3KCA* variants, has an average log(IC_50_) of 1.01µM for selumetinib and −0.672µM for afatinib. In contrast, the *BRAF* mutation was enriched in a subpopulation where the average log(IC_50_) for selumetinib was −1.097µM and 0.625µM for afatinib. The difference in response between afatinib and selumetinib was significantly greater (t-test P<0.01) between the subpopulations identified and the total population of *PIK3CA* or *BRAF* mutant cell lines (**Figures 1F and 1G**).

### Cross-comparison of multiple drugs redefines best-in-class drugs for specific subpopulations

Although there is a larger portfolio of clinical drugs with identical putative targets, their responses may differ substantially in subpopulations as a consequence of multiple factors, for example mode-of-action, different off-target effects and binding properties. The ability to discover cell line subpopulations with distinct pharmacological patterns of response characterised by genetic mutations re-defines best-in-class drugs by their differential response to other drugs in a specific subpopulation, rather than their absolute response across an entire population.

In order to demonstrate this approach for drug discovery, we applied SEABED to 745 cell lines across cancer types to evaluate the differential response in those cell lines to five inhibitors (CI-1040, PD0325901, RDEA119, selumetinib, and trametinib) which all target the MEK protein (**Figure 2A**). The segmentation of cell lines revealed 13 subpopulations with different patterns of response and three having enriched biomarkers (**Figure S2A**). Two subpopulations were sensitive to all MEK inhibitors, with trametinib achieving the greatest sensitivity. In one subpopulation the *KRAS* mutation was enriched (Fisher exact p-value = 1.12e-4 and 40.8% of the cell lines) while another had the *BRAF* mutation enriched (Fisher exact p-value = 1.39e-7 and 50% of the cell lines). In contrast, another subpopulation was enriched with the *RB1* mutation (Fisher exact p-value = 3.84e-2 and 21.6% of cell lines), within which the cell lines were almost uniformly resistant to all MEK inhibitors.

**Figure 2:**
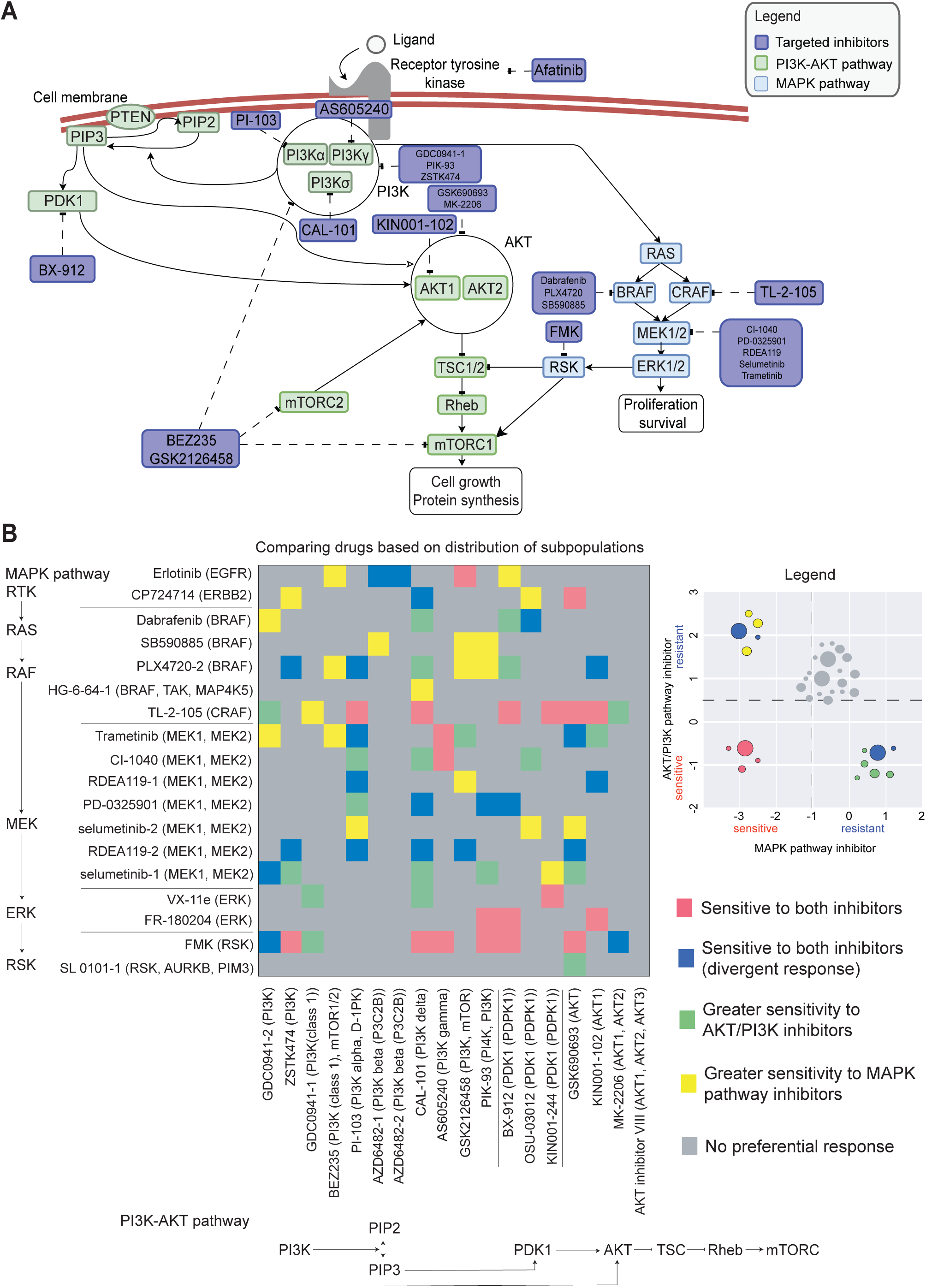
Distinct drug response types after unsupervised segmentation of pharmacological response pattern for targeting MAPK or PI3K-AKT signaling. **(A)** MAPK and PI3K-AKT pathways illustrating drugs in purple boxes which were assessed, and their different gene targets in the pathway. Genes in the green boxes are involved in the PI3K-AKT pathway while genes in the blue boxes are involved in the MAPK pathway. **(B)** Heatmap illustrating 324 pairwise comparison of responses for 18 different inhibitors targeting the MAPK pathway and 18 different inhibitors targeting the PI3K-AKT pathway. The 20th-percentile of log(IC_50_) values for each drug was determined based on the distribution of log(IC_50_) values across all cell lines after SEABED segmentation. Based on these 20th percentile cutoffs for sensitivity, we assessed whether there was enrichment of subpopulations in each quadrant and categorised each drug pair into five categories: (i) no preferential response (grey), (ii) subpopulations sensitive to both MAPK and PI3K-AKT pathway inhibitors (pink), (iii) greater MAPK pathway sensitivity (yellow), (iv) greater PI3K-AKT pathway sensitivity (green) or (v) sensitive to either inhibitor but not both (blue).

### Distribution of subpopulations highlight distinct pharmacological relationships between PI3K-AKT and MAPK signaling

Next, we used SEABED to investigate the cross-talk between two frequently active cancer pathways, MAPK and PI3K-AKT signalling, by systematically comparing pairs of drugs targeting different genes of each pathway (**Figure 2A, B**). In total, SEABED performed 324 pairwise comparisons of 18 PI3K-AKT and 18 MAPK pathway inhibitors. Each drug pair was classified into five categories based on the distribution of subpopulation responses: (i) no preferential response, (ii) sensitive to both MAPK and PI3K-AKT pathway inhibitors (**Figure S2B**), (iii) greater sensitivity to MAPK pathway inhibitors (**Figure S2C**), (iv) greater sensitivity to PI3K-AKT pathway inhibitors (**Figure S2D**), (v) sensitive to either a MAPK pathway or a PI3K-AKT pathway inhibitor, i.e. divergent response (**Figure S2E**). While this approach classified the distribution of subpopulations based on somewhat arbitrary thresholds for sensitivity, we also measured the weighted average Pearson correlation across subpopulations to identify interesting drug pairs (**Figure S3**).

We found 20 drug pairs subpopulations with sensitivity to both PI3K-AKT and MAPK pathway inhibition. This association between subpopulation size and sensitive response was significant when comparing a CRAF inhibitor (TL-2-105) to PI3K-AKT signaling inhibitors (hypergeometric test P=2.358e-5). The same trend was observed for inhibiting ERK (FR-180204) or RSK (FMK) compared to inhibiting any PI3K-AKT signaling gene (P=0.0185 and P=2.358e-5, respectively; hypergeometric test), but interestingly there was no mutual sensitivity when comparing to BRAF inhibitors and only two cases for MEK inhibitors.

There were 20 drug pairs with a significantly high proportion of subpopulations (P < 0.05) exhibiting greater sensitivity to MAPK pathway inhibition. This phenotype is strongly pronounced in pairs with BRAF inhibitors (hypergeometric test P=0.0166). 20 drug pairs were found with significantly high proportions of preferential PI3K-AKT pathway inhibition. This is observed when comparing MEK inhibitors (CI-1040 and selumetinib-1) to PI3K-AKT signaling inhibitors (P=0.0185 and P=0.00273 respectively; hypergeometric test).

In 22 cases, we observed drug pairs with sensitivity to either a MAPK pathway or a PI3K-AKT pathway inhibitor, i.e. divergent response. This response type was enriched for pairs of any PI3K-AKT pathway inhibitors and BRAF (PLX4720-2) or MEK inhibitors (P=0.026 and P=0.0135, respectively; hypergeometric test), and even significant for PI3K inhibitors in comparison with either the EGFR (erlotinib) or MEK inhibitors (P=0.0315 and P=0.0173; hypergeometric test). Response patterns for all drug pairs can be explored in our portal (Website S1; https://szen95.github.io/SEABED).

### Subpopulations of differential response identifies drug combination efficacy

Previous studies have hypothesised that the efficacy of many approved drug combinations can be explained by the independent action of single agents on different patient subpopulations with cancers driven by multiple pathways ^21^. We hypothesised that SEABED comparisons of drug pairs would highlight subpopulations of differential response that would exhibit synergistic or independent action effects when the drugs are tested in combination. Additionally, our method enables to explore putative biomarkers of such populations. Systematic comparison of responses between two drugs highlighted subpopulations of cell lines in which there was sensitivity to either drug but not both (divergent response). We observed this phenomenon in 22 drug pair comparisons, including a MEK inhibitor (RDEA119-2) which showed divergent responses to four PI3K inhibitors (PI-103, GSK2126458, ZSTK474, and CAL-101; **Figure 3A; Figures S4A-D** respectively). Drug pairs with divergent response were also observed in cell lines treated with PLX4720-2 (BRAF inhibitor) and two PI3K inhibitors (PI-103 and ZSTK474; **Figures 3B and S4E; Figure S4F**). Two subpopulations with a high proportion of *BRAF* mutations were identified with greater sensitivity to the BRAF inhibitor (**Figure 3B**). These two subpopulations had a combined total of 79 cell lines (**Figure 3C**). The subpopulation with an average log(IC_50_) of −0.0232µM for PLX4720-2 and 0.771µM for PI-103 had 21 cell lines with *BRAF* mutation (87.5%; P=9.75e-17). In contrast, the subpopulation with an average log(IC_50_) of 0.808µM for PLX4720-2 and 0.562µM for PI-103 had 22 cell lines with *BRAF* mutation (40.4%; P=5.12e-(**Figure 3B**). We observed 17 individual cell lines with *BRAF* mutation that are resistant to both drugs (**Figure S4G**).

We next examined the drug pairs as combination therapies in cell lines ^25^ and patient-derived tumor xenograft models (PDXs) ^26^ to investigate whether the drug pairs with divergent response and subpopulations with preferential sensitivity to one drug would be associated with efficacy of their combination treatment (**Figure 3D**). SEABED first compared the single drug responses of BRAF, MEK and PI3K inhibitors as before to identify BRAF mutant subpopulations with differential response. When the drugs were tested as combinations in BRAF mutant cell lines, the MEK/PI3K inhibitor combination had a surprising similar level of synergy as BRAF/MEK combinations, which was recently a clinically approved combination ^27,28^. Also surprising, these two combinations had significantly higher synergistic effect when used on BRAF mutant cell lines compared to all cell lines (t-test P=0.0204), and compared to all drug combinations tested (t-test P=1.46e-5; **Figure 3E; Figure S4H**). In terms of overall efficacy in PDXs, we observed a similar level of inhibition to tumour volume for the BRAF/PI3K inhibitor combination on BRAF mutant cells when compared to the clinically approved BRAF/MEK combination and a significantly greater (t-test P=0.0418) inhibition of tumour growth compared to all combinations (**Figure 3F; Figure S4I**). Notably, previous work suggested *in vivo* efficacy of drug combinations is mostly driven by the monotherapy agents targeting independent mechanisms ^26^; however, they did not exclude the possibility that such drug combinations may also be synergistic. Here, we highlight a drug combination example, where *in vivo* efficacy is driven by targeting independent mechanisms, and complementary being synergistic. This example highlights that both concepts, synergy and targeting independent mechanism may contribute to combination efficacy in patients.

**Figure 3:**
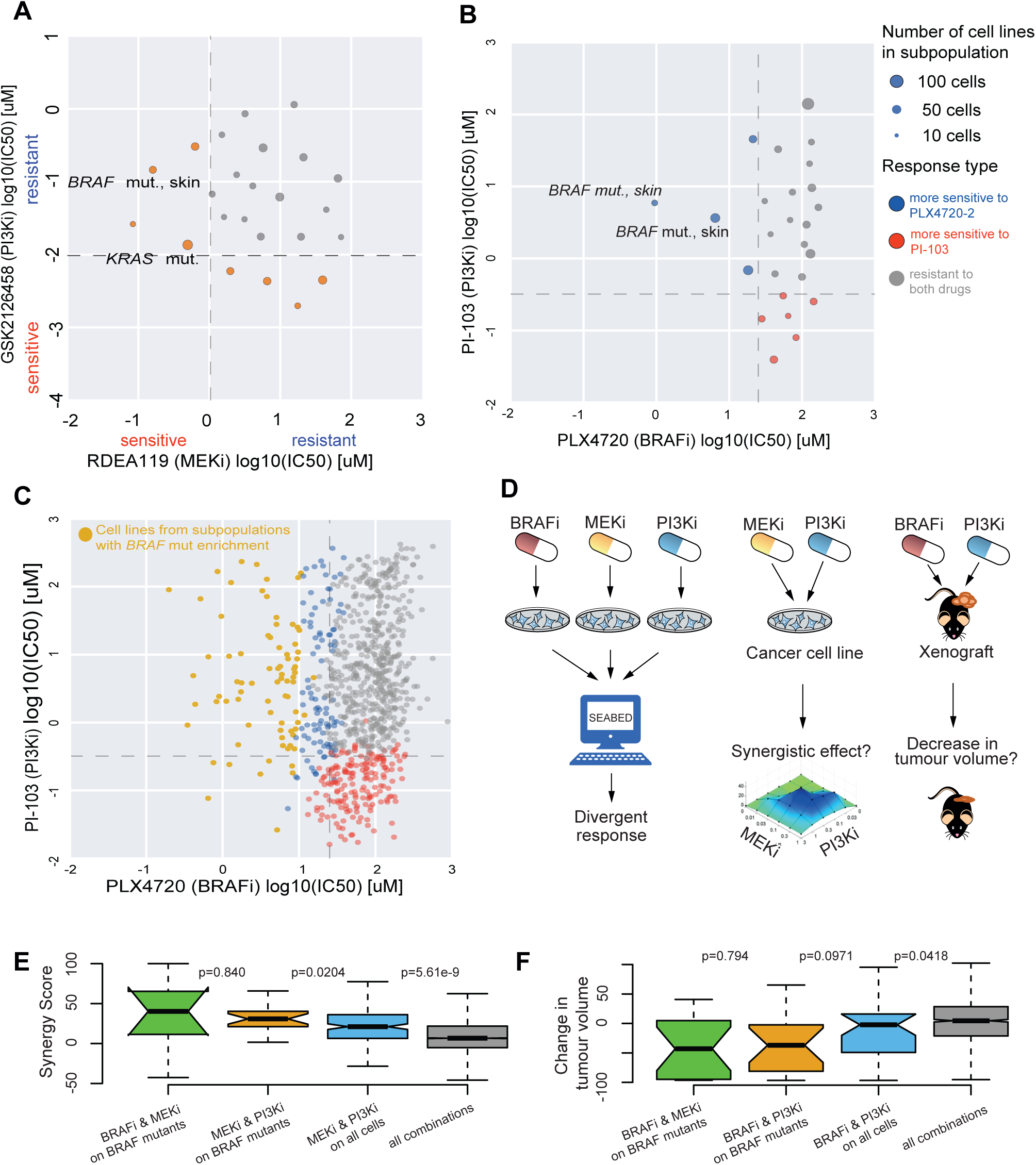
Divergent response exemplified with PI3K inhibition in comparison to MEK and BRAF inhibitors. **(A)** Scatter plot showing subpopulations that exhibit divergent pharmacological response to RDEA119-2 (MEK inhibitor) and GSK2126458 (PI3K inhibitor). Dashed lines indicate 20th percentile of log(IC_50_) values for each drug. *BRAF* and *KRAS* mutations were found enriched in the associated subpopulations. **(B)** Subpopulations from comparison of PLX4720-2 (BRAF inhibitor) and PI-103 (PI3K inhibitor) responses. **(C)** Individual cell line responses from PLX4720-2 and PI-103 coloured by the subpopulations they were grouped in. **(D)** Workflow illustrating cell lines being tested with individual inhibitors and their joint pharmacological patterns of response. Drug pairs with divergent response and *BRAF* mutant subpopulations suggest a target for drug combination therapies, which are validated in cell lines ^25^ and patient-derived tumor xenograft (PDX) models ^26^. **(E)** *In vitro* synergistic effect of combining MEK inhibitors with PI3K or BRAF inhibitors in all cell lines and just those with *BRAF* mutations. This also compared to measured synergy for all combinations tested in cell lines. **(F)** *In vivo* effect of combining BRAF inhibitors with PI3K or MEK inhibitors. Measured response is the change in tumor volume following treatment. BRAF and MEK inhibitor combinations have also been shown to be effective in the patients ^28^.

### Lack of subpopulations of differential response may explain clinical failure

Sometimes, despite strong preclinical evidence, some drugs do not succeed in clinical trials ^29^. One such trial was SELECT-1 (**Table S1**) which compared the efficacy of combining selumetinib and docetaxel to docetaxel alone in patients with advanced *KRAS*-mutant non– small cell lung cancer (NSCLC) ^30^. Although there were *KRAS* mutant cell lines sensitive to selumetinib in preclinical testing ^31^, we re-examined the pharmacological data with SEABED to assess whether there were distinct subpopulations that justified the patient selection criteria for *KRAS* mutation.

In this analysis, instead of only inspecting the subpopulation identified by SEABED when the segmentation algorithm terminated, we thoroughly examined all possible subpopulations. SEABED identified a total of 61 possible subpopulations from 840 cell lines across tissue types tested with selumetinib and docetaxel (**Figure 4A**). 10 subpopulations were more sensitive to selumetinib than docetaxel (**Figure 4B**), and 5 of those subpopulations were enriched for *KRAS* mutation. However, those subpopulations enriched for NSCLC *KRAS* mutants were small in size and mostly exhibited less sensitivity to selumetinib compared to docetaxel (**Figures S5A and S5B**). The distribution of different *KRAS* mutations (p.G12C vs p.G12V) was also no different in selumetinib sensitive subpopulations compared to resistant subpopulations (**Figures S5C and S5D**). Independent of mutation status, only 8.9% of NSCLC cell lines were found in subpopulations more sensitive to selumetinib, whereas 14.8% cell lines originating from aerodigestive cancer types (eg. esophageal) were found in these subpopulations (**Figures S5E and S5F**).

**Figure 4:**
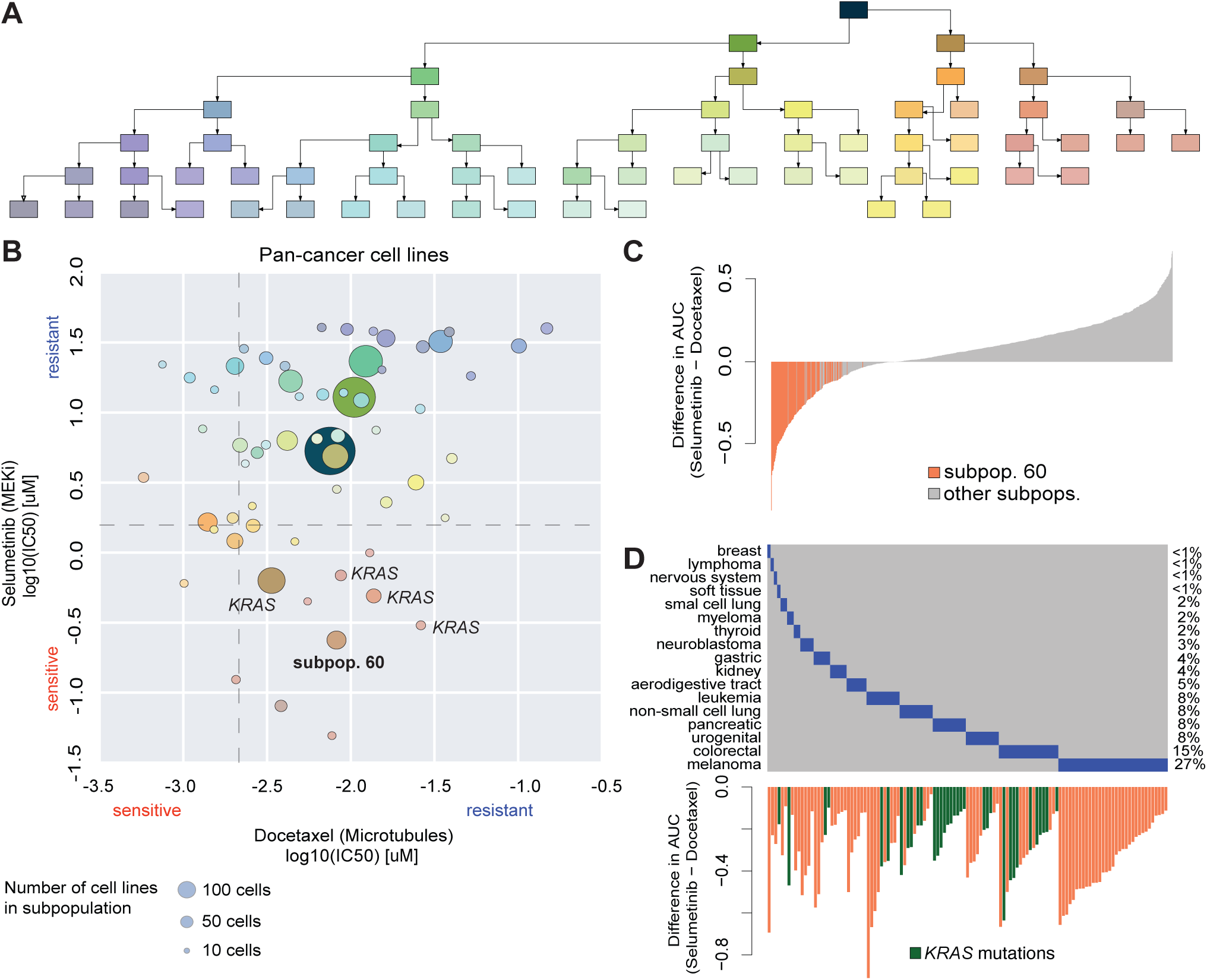
Subpopulations of cell lines indicating differential response between docetaxel and selumetinib. **(A)** Tree diagram illustrating the segmentation process of 840 cell lines across cancer types into subpopulations based on their response to docetaxel and selumetinib. Branch colours distinguish subsets of subpopulations with darker colours indicating increasing number of cell lines in the subpopulation. **(B)** Scatter plot of subpopulations discovered through progressive segmentation based on average log(IC_50_) values. Dashed lines indicate 20th percentile of log(IC_50_) values for each drug. The *KRAS* mutation is enriched in 5 subpopulations exhibiting sensitivity to selumetinib and resistance to docetaxel, including subpopulation 60. The colours of the subpopulations correspond to the location of the subpopulation in the tree diagram. **(C)** Bar plot illustrating the difference in AUC values for each cell line. The orange bars highlight the cell lines within subpopulation 60. **(D)** Heatmap showing the percentage of each cancer type enriched within the cell lines in subpopulation 60. The bar plot illustrates the *KRAS* mutations within the cell lines that are highlighted in green.

Next, we focused on *subpopulation_60*, which had the greatest difference in sensitivity (IC_50_ and AUC) to selumetinib compared to docetaxel (**Figure 4C**). This subpopulation of 122 cell lines was enriched in *KRAS* mutations (28.8%, P=3.061e-4) found across multiple tissue types. NSCLC cell lines accounted for only 8% of this subpopulation, with 50% of those cell lines being *KRAS* mutants. Colorectal and pancreatic cell lines accounted for 15% and 8% respectively of the subpopulation, and they both had a higher proportion of *KRAS* mutations (56% and 100% respectively; **Figure 4D**).

## Discussion

The ability to identify distinct subpopulations based on multiple measures of drug response (eg. IC_50_ and AUC) and extract their biomarkers is the basis for personalised therapeutics, which may ultimately increase the likelihood of successful clinical trials ^32,33^. Using a network-based segmentation algorithm coupled with biomarker detection (SEABED), we investigated well-established pharmacological targets and clinical biomarkers by comparing the response patterns for BRAF (SB590885) and MEK (CI-1040) inhibition, which expectedly reproduced subpopulations sensitive to both enriched for *BRAF* mutants ^34–36^. In another example, SEABED compared EGFR/ERBB2 (afatinib) and MEK (selumetinib) inhibition to reveal expected biomarkers such as *BRAF, KRAS* and *NRAS* mutations for selumetinib ^13–16^, and afatinib associated with *EGFR* and *ERBB2* amplifications ^37,38^. Interestingly, the more afatinib-resistant subpopulation was enriched for *PI3KCA*-activating mutation, which may cause acquired resistance ^24^. When we systematically compared inhibitors of the MAPK and PI3K-AKT signaling pathways, we observed subpopulations sensitive to both CRAF, ERK or RSK targeted drugs and other drugs targeting the PI3K-AKT pathway, however, there were few instances of these subpopulations for inhibitors targeting other genes in the MAPK signalling ^*39*^. We found many more subpopulations that were more sensitive to BRAF inhibitors than other PI3K-AKT inhibitors, and as expected, many contained *BRAF* mutations ^34^. In contrast, there were not significantly more subpopulations sensitive to MEK inhibition compared to inhibition of PI3K-AKT signalling targets, but *BRAF* mutant subpopulations may have greater differential response ^14^. Divergent response was observed when comparing EGFR, BRAF and MEK inhibitors to drugs targeting the PI3K-AKT pathway. Our results comparing the MAPK and PI3K-AKT pathways based on drug response profiles highlights how intertwined those two pathways are in pharmacology space ^39^.

Arguably, the divergent response type is the most exciting for personalised treatment, since it may identify cases where independent drug action and synergy may guide effective drug combinations ^21^. Here exemplified, we showed that PI3K inhibitors combined with either BRAF or MEK inhibitors increase *in vitro* synergy and reduce tumour volume of *in-vivo* models. Furthermore, we were able to show that synergistic and overall effect can be further enhanced by the correct biomarker indication, in this instance, *BRAF* mutant subpopulations ^40,41^. The *BRAF* mutant subpopulation with high efficacy for the BRAF inhibitor and not the other inhibitor could be cases where independent drug action explains drug combination efficacy, whereas, the subpopulation with lower efficacy for single treatments of either drug may be cases for synergistic effects when the drugs are combined.

In examining the preclinical evidence for trial testing combination treatment of NSCLC in which the *KRAS* mutation was the biomarker ^42^, SEABED revealed a high proportion of NSCLC subpopulations having the *KRAS* mutation that are resistant to both selumetinib and docetaxel, suggesting a smaller likelihood of efficacy for the drug combination. Alternately, we identified a subpopulation with differential response to selumetinib for a small proportion of *KRAS* NSCLC cell lines, but this subpopulation contained a higher proportion of colorectal and pancreatic cancer cells with *KRAS* mutations. Previous studies have shown the plausibility in treating colorectal cancer using MEK inhibitor combinations ^43,44^. With consideration of *KRAS* mutations in subpopulations having greater sensitivity to selumetinib, SEABED suggests that while the correct biomarker was used for the clinical trial, there may be other potential indications for selumetinib. Although response in cell lines may not always correspond to response clinically, the use of data-informed approaches to examine large populations of cells may reveal clinically relevant drug response patterns. Future studies may need to account for differences between *in vitro* and *in vivo* responses.

While iterative and hierarchical clustering techniques have been used widely to attribute molecular markers to differences in subpopulation drug response and outcomes ^45,46^, we use an approach that does not require an explicit estimate of the number of subpopulations and is not greedy, i.e., each incremental step is optimal but the overall algorithm is not. In biomedical data processing, there has been substantial concern, particularly regarding applications to molecular data, that rival unsupervised machine learning optimize different criteria and consequently yield diverging answers ^47,48^. In our effort, we are concerned with discovering subpopulations having high homogeneity and statistical separability, while avoiding subpopulations that are so small that extracting statistically significant biomarkers is unlikely. We demonstrated the utility of SEABED over conventional approaches of K-Means and hierarchical clustering for drug response comparisons (see **Supplementary Materials, Table S2, Table S3, and Figure S6**). Further, the ability to segment non-convex regions, which can arise to describe different disease states, is advantageous. Consequently, because of their success in other industries ^49,50^ as well as fast, efficient implementations, spectral methods based on network models are powerful methods for discovering distinct subpopulations ^51–53^.

SEABED builds upon previous work using network models in biomedical contexts ^51,54,55^ that explicitly partition a population of cell lines described by multiple variables into distinct subpopulations, using a “top-down” approach of recursively identifying optimal cuts for graph bisection. Similarly, while our segmentation capitalizes on past progress made in spectral clustering ^56,57^, our effort distinguishes itself from past attempts by integrating all variables into a single network model using a multivariate similarity and local and global network statistics. Deeper interpretations of embedded matrix subspaces in network models may provide further insight into the linkage between subpopulations of cancer cell lines and drugs.

As a whole, this study demonstrates several important insights about the pharmacological pattern of response for different cancer drugs by applying an unsupervised machine learning platform to segment a large pan-cancer *in vitro* pharmacology data set. By organizing cell lines along similar pharmacological patterns of response, we identified distinct, intrinsic subpopulations sensitive to one drug but resistant to others, and in some cases identified genetic alterations that can be used as biomarkers for those subpopulations. In the context of analytical frameworks for increasing drug R&D productivity by sharpening the focus of drugs ^58^, our work demonstrates the value of advanced analytical approaches in translational medicine to enable decision making that is more data-informed and less ambiguous. Moreover, by analyzing different pharmacological responses and interpreting its outputs in the context of the underlying genetics and molecular pathways, we have created a multifaceted landscape for developing and assessing new drug therapies.

## Methods

### CONTACT FOR REAGENT AND RESOURCE SHARING

All code for the pipeline is open source and available at: https://github.com/szen95/SEABED. Further information and requests should be directed to and will be fulfilled by the co-corresponding authors, Michael P. Menden or Dennis Wang.

### METHOD DETAILS

#### Pharmacology data

The discovery pharmacology dataset was extracted from the The Genomics of Drug Sensitivity in Cancer (GDSC) database ^3,4^, while leads from the analysis were validated with the Cancer Cell Line Encyclopedia (CCLE) ^5^ and the Cancer Therapeutics Response Portal (CTRP) ^6–8^. Furthermore, suggested drug combinations were validated with cell line responses from the AstraZeneca-DREAM challenge dataset ^25^ and patient derived xenograft (PDX) models from Gao *et al.* ^26^.

For a given cell line in GDSC, the drug response was fitted with a sigmoid curve ^59^ and consecutively quantified as area under the curve (AUC) or the concentration required to reduce cell viability by half (IC_50_). GDSC contains 265 compounds tested in 1074 cell lines, whilst we focus on a subset of 38 drugs targeting either the PI3K-AKT or MAPK signalling, which leads to 327 experiments considered for evaluation.

#### Deep molecular characterisation of the cancer cell lines

The GDSC project ^4^ provides the characterisation of >1,000 cell lines including whole exome sequencing, targeted PCR sequencing/split probe FISH analysis and SNP6.0 arrays, which enabled to quantify somatic mutations, gene fusions and copy number variations (CNVs), respectively. In our analysis, we focus on somatic mutational state of 300 cancer genes and 10 gene fusions. Additionally, we considered 425 recurrent CNVs, split into 117 amplifications and 308 deletions. In total, we consider 735 cancer functional events, which is summarized in the binary event matrix (BEM) from Iorio *et al.* ^*4*^.

#### Pre-processing cell line and pharmacological data

For every pair of drugs that was computationally analyzed, the subset of GDSC cell lines having valid IC_50_ and AUC values for both drugs was retained. Typically there were roughly 700 cell lines across all cancer types that met these criteria in each experiment. Cance types and subtypes were stored along with the pharmacological data in a table for each drug for subsequent recall and analysis.

#### Processing drug response measures (AUC/IC_50_ values)

We build network models for a set of *N* cell lines, *C* = {*C*_1_,.. *C*_*N*_}, that are separately exposed to two distinct drugs, *D*_1_ and *D*_2_, which results in two sets of *M* measurement variables, *X*_*i*_ = [*x*_1_,.., *x*_*M*_], *i* = 1,2, describing the response to each compound:

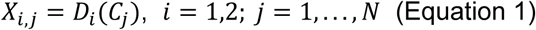

We use a network model that is an undirected graph, *G*, consisting of *N* vertices, *V*_*i*_,*i* = 1,…, *N*, (one for each cell line in *C*) with weighted edges, *W*_*i,j*_(*V*_*i*_, *V*_*j*_), *i,j* = 1,…, *N,i* ≠ *j*, between every distinct pair of vertices. Our approach uses a single multivariate similarity measure (Equation 2), to construct one network model, with the advantage that the subspace properties of the resulting adjacency and Laplacian matrices are fully embedded with the complete characteristics of *C*. The weight is the similarity, w_i,j_, between *i*-th and *j*-th composite *2M x* 1 dose response profile (DRP), *X*_*i*_ = [*X*_1,*i*_, *X*_2,*i*_], for *C*_*i*_ and *C*_*j*_.

We characterize drug response by two important continuous-valued measurements extrapolated from the cell line pharmacology screens: the IC_50_ and the AUC values of the dose-response curve (**Table S4**) observed when one compound is applied *in vitro* to a single cell line sample at successively greater concentrations. Since every cell line possesses a length-4 DRP for a given pair of drugs, the similarity, *W*, between any two cell lines resides on (0,1) and is calculated by a multivariate quasi-Gaussian comparison that differences the elements of the DRPs but also weighs the differences by a combination of local and global network statistics. Similarity between the response vectors, *X*_*i*_ and *X*_*j*_, is given by:

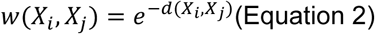

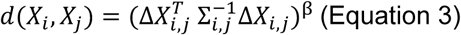

The similarity between two cell lines equals one when both have identical covariate values, and approaches zero as their covariates increasingly differ. Additionally, *w*(*X*_*i*_, *X*_*j*_)=*w*(*X*_*j*_, *X*_*i*_). Δ*X*_*i,j*_ is a *4 ×* 1 vector whose entries are the difference of the DRP values in *X*_*i*_ and *X*_*j*_ and β modulates the similarity between two patients. We selected β = 0.5 for our experiments based on experimentation and the observations of previous efforts.

Σ_*i,j*_ is a 4 × 4 covariance-like matrix that is estimated for every distinct (*i, j*)-pair and captures the variability of individual variables as well as their inter-relationships. While Σ_*i,j*_ is an explicit function of the two patients being compared, it also captures network-wide characteristics. For diagonal elements, Σ_*i,j*_(*a, a*), *a* = 1,..4, the entries are:

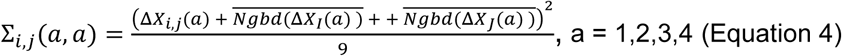

where *Ngbd* (Δ*X*_*I*_(*a*)) corresponds to all edges neighboring vertex-i, and the overbar is the averaging operator. The off-diagonal elements, Σ_*i,j*_(*a, b*), *a, b* = 1,..4, *a* ≠ *b* are:

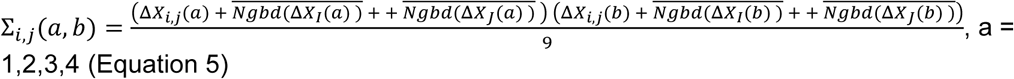

The Moore-Penrose pseudo-inverse was used to avoid problems with low-rank during matrix inversion. The framework is generalizable to include more variables and different measures of similarity. The symmeric, positive semi-definite, *N x N* weighted adjacency matrix, *W*, holds the pairwise similarities.

#### Segmentation

The set of cell lines, *C*, is segmented recursively into distinct subpopulations using the Fiedler eigenvector derived from the eigendecomposition of *W* ^60^. Each parent subpopulation of cell lines is successively segmented into two offspring subpopulations until at least one of 3 constraints is satisfied:

- The size of the parent subpopulation falls below a user-defined threshold;
- The size of either offspring subpopulation is below a user-defined threshold;
- The offspring subpopulations are not sufficiently dissimilar, where we measure class separability using the silhouette metric ^61^, a common non-parametric method in which values range between −1 (highly similar) and 1 (highly dissimilar) and values near 0 indicate the two subpopulations are just barely overlapping. Higher values result in fewer, less granular, subpopulations.

In our experiments, we required the size of the parent subpopulation to be at least 40 in order for segmentation to be performed, and required both offspring subpopulations to have 20 or more members in order to be retained. The silhouette metric threshold was set to 0.25. Generally, criteria and thresholds can be modified and adapted to emphasize relevant factors in a particular problem. The whole segmentation process yields *K* mutually exclusive subpopulations *P*_*k*_, *k* = 1,…, *K*, where *C = U*_*k*=1_ *P*_*k*_. Successive segmentation results in sub-populations with increasingly homogeneous DRPs.

### QUANTIFICATION AND STATISTICAL ANALYSIS

#### Enrichment of features to nominate biomarkers

Because genetic alterations in each cell line are known, each subpopulation can be evaluated by non-parametric statistical tests to identify enriched alterations that may be attributed to patterns of sensitivity or resistance in the DRP across both drugs. For each subpopulation, we measured the number of cell lines in the subpopulation with a particular gene mutation, and the number of cell lines outside of the subpopulation with the mutation. A 2×2 contingency table was generated from the cell line counts of with/without mutation and inside/outside of subpopulation. Significance of observed enrichment of mutations within subpopulations were calculated using the Fisher’s exact test. The resulting p-values were corrected for multiple testing using the Benjamini and Hochberg (BH) procedure (**Table S5**).

#### Classification of drug pairs based on the distribution of subpopulations

We made 324 pairwise comparisons of drugs targeting the MAPK and PI3K-AKT pathways. Based on the distribution of log(IC_50_) values across all cell lines tested with both drugs, we determined the 20th-percentile of log(IC_50_) values for each drug. The 20th percentile cutoffs *P*_*20*_ for drugs A and B was used to categorise each subpopulation *i* into four categories based on their average log(IC_50_) *y*:

*y*_*i*_ *< P*_*20, A*_ and *y*_*i*_ *< P*_*20, B*_ *= sensitive to drugs A and B*

*y*_*i*_ *< P*_*20, A*_ and *y*_*i*_ ≥ *P*_*20, B*_ *= more sensitive to drug A*

*y*_*i*_ ≥ *P*_*20, A*_ and *y*_*i*_ *< P*_*20, B*_ *= more sensitive to drug*

*y*_*i*_ ≥ *P*_*20, A*_ and *y*_*i*_ ≥ *P*_*20, B*_ *= resistant to drugs A and B*

The number of subpopulations in each category were recorded in a 2×2 contingency matrix and using a binomial test compared this to the proportion of cells expected in each category if SEABED segmentation was not performed.

After classification of pairwise drug responses, we assessed whether a drug was significantly enriched for one category in comparison with all other drugs. Testing was carried out using the hypergeometric test (*phyper* R package).

#### 2-D visualization of drug response profiles

To visualize DRPs across cell lines and drug comparisons, we calculated the average log(IC_50_) values for each drug in subpopulations generated based on their response to the tested drug pairs (**Table S5**). We then plotted the mean log(IC_50_) values as circles on a 2-D scatter plot using the Matplotlib Python library. Dashed lines indicative of local 20th percentile of log(IC_50_) values for each drug were also plotted on the scatter plot unless stated otherwise. The radii of the circles is proportional to the subpopulation size. Due to potential misidentification, cell lines named ‘NA’ found in the output files after segmentation with SEABED, were not used for the calculation of the mean log(IC_50_) values of each subpopulation and hence, not used for the 2-D visualization.

#### Tree visualization of subpopulations

We utilized tree diagrams to visualize the data generated. The tree diagrams illustrate how the cancer cell lines are segmented into different subpopulations, based on whether they are sensitive or resistant to the drugs that are being tested. We did not visualize further down the tree in the figures when previously observed significant enrichment of genetic biomarkers is no longer observed in all current subpopulations. The tree diagrams were generated through an open-source Python library called Graphviz. The style of each component of the tree diagram was first initialized through a class. This included the colours, shapes, and fonts of the edges and nodes of the tree diagram. A method to create tree diagrams was developed to accept the number of vertices and leaves, the labels for the leaves, and the tree diagram filename. The tree diagram is finally generated and saved by calling the method.

### DATA AND SOFTWARE AVAILABILITY

All code for the pipeline is open source and available at: https://github.com/szen95/SEABED. All data used in the paper are published previously and publicly available at the GDSC, CCLE, and CTRP databases. Datasets used are listed in Table S4, Table S5, and the Key Resources Table.

### ADDITIONAL RESOURCES

Response patterns of 324 pair-wise comparisons of 18 PI3K-AKT and 18 MAPK pathway inhibitors: https://szen95.github.io/SEABED/

## Supplemental Information

1. Supplementary Materials.pdf Document S1. Figures S1–S6 and Tables S1-3.
2. Table S4.xlsx Table S4. Input Data for Segmentation, Related to Figures 1-4, S1-S6, and Website S1 **Input data:** log(IC_50_) and AUC values for 1074 cancer cell lines treated with 265 anti-cancer drugs.
3. Table S5.xlsx Table S5. Output Data and Enriched Biomarkers After Segmentation, Related to Figures 1-4, S2-S6, and Website S1 **Output data:** IC_50_ 20% cutoff, minimum and maximum IC_50_ concentration, and average IC_50_ response of each subpopulation towards each drug (327 cancer drug pairs) tested on the cancer cell lines. The subpopulation number and the number of cell lines in each subpopulation are recorded. Each individual cell line in every subpopulation together with individual cell line tissue types are also shown. **Enriched biomarkers:** Biomarkers found within subpopulations (adjusted p-value and/or p-value < 0.05), together with the subpopulation number, the number of cell lines in each subpopulation, percentage of the biomarkers found within each subpopulation, the number of cell lines in the subpopulation with the biomarker, the p-value, and adjusted p-value.

## Acknowledgements

We would like to thank Ben Sidders (AstraZeneca plc.), Jonathan Dry (AstraZeneca plc.), Francesco Iorio, (Sanger Institute), Michael Schubert (EMBL-EBI), Mi Yang (RWTH Aachen) and Winston Hide (Harvard University) for useful discussions. This work is supported by funding from the NIHR Sheffield Biomedical Research Centre, Rosetree Trust (ref: A2501), and the Academy of Medical Sciences Springboard (REF: SBF004\1052).

## Author contributions

NK, TST, MM, and DW contributed to the conceptualization of the project. NK, TST, and DW were responsible for data curation. Formal analysis was performed by NK, TST, HY, BY, MM, and DW. MM and DW were responsible for funding acquisition. The methodology was developed by NK, TST, MM, and DW. TST and DW were responsible for administration of the project. Resources for the experiments were prepared by DW and NK. Various software for the project was developed and implemented by NK, TST, HY and DW. MM, and DW oversaw supervision of the project. NK, TST, MM, and DW wrote the manuscript.

