## Supplementary Materials for "Defining subpopulations of differential drug response to reveal novel target populations"

### Contents

**Comparison to K-Means and hierarchical clustering 2**
**Supplementary Figures 3**
**Supplementary Tables 10**
**Additional References 12**

#### Comparison of SEABED Segmentation Results with Existing Methods

To yield meaningful and usable subpopulations, we considered the performance of SEABED with respect to hierarchical clustering (Euclidean Minimum Distance and single-linkage) and K-Means clustering (Euclidean Minimum Distance). To enable equitable comparisons, both methods were constrained to find the identical number of subpopulations as SEABED. We were primarily concerned with factors impacting segmentation such as selecting the overall dimensionality of the decomposition, the aggregate separability of the subpopulations, and the size of the subpopulations. The latter two factors are especially important since they directly impact the ability to reliably extract distinct biomarkers.

Two compounds, PLX4720-2 (BRAF inhibitor) and PI-103, were examined in Figure 3 using SEABED. Using the same number of subpopulations found by SEABED, 25, we also executed hierarchical clustering and K-Means to also be evaluated for 25 subpopulations. Our results in Table S2 demonstrate that SEABED identifies the most balanced subpopulation sizes, with the largest minimum and median subpopulation size, which is conducive to finding biomarkers. Similarly, we evaluated two different cluster validation measures to evaluate the uniqueness of each pair of subpopulations. Our results in Table S3 demonstrate that SEABED finds competitive separation values for the silhouette metric and also has the highest subpopulation homogeneity (RMSSTD). Furthermore, SEABED did not require a prior definition of the number of subpopulations, which is a mandatory input for K-Means.

Finally, we investigated what biomarkers were found by K-Means and hierarchical clustering for the drug pair in Figure 3B (PLX4720-2 (BRAF inhibitor) and PI-103). In Figure S6, the results for SEABED are compared to the two competing methods. Hierarchical clustering produced one large cluster and several singleton results, making biomarker discovery in sensitive regions impractical. K-Means clustering found two subpopulations enriched for *BRAF* mutations, but with higher P-values. For K-Means clustering, the divergent region had more subpopulations, but they were substantially smaller than the ones found by SEABED.

### Supplementary Figures

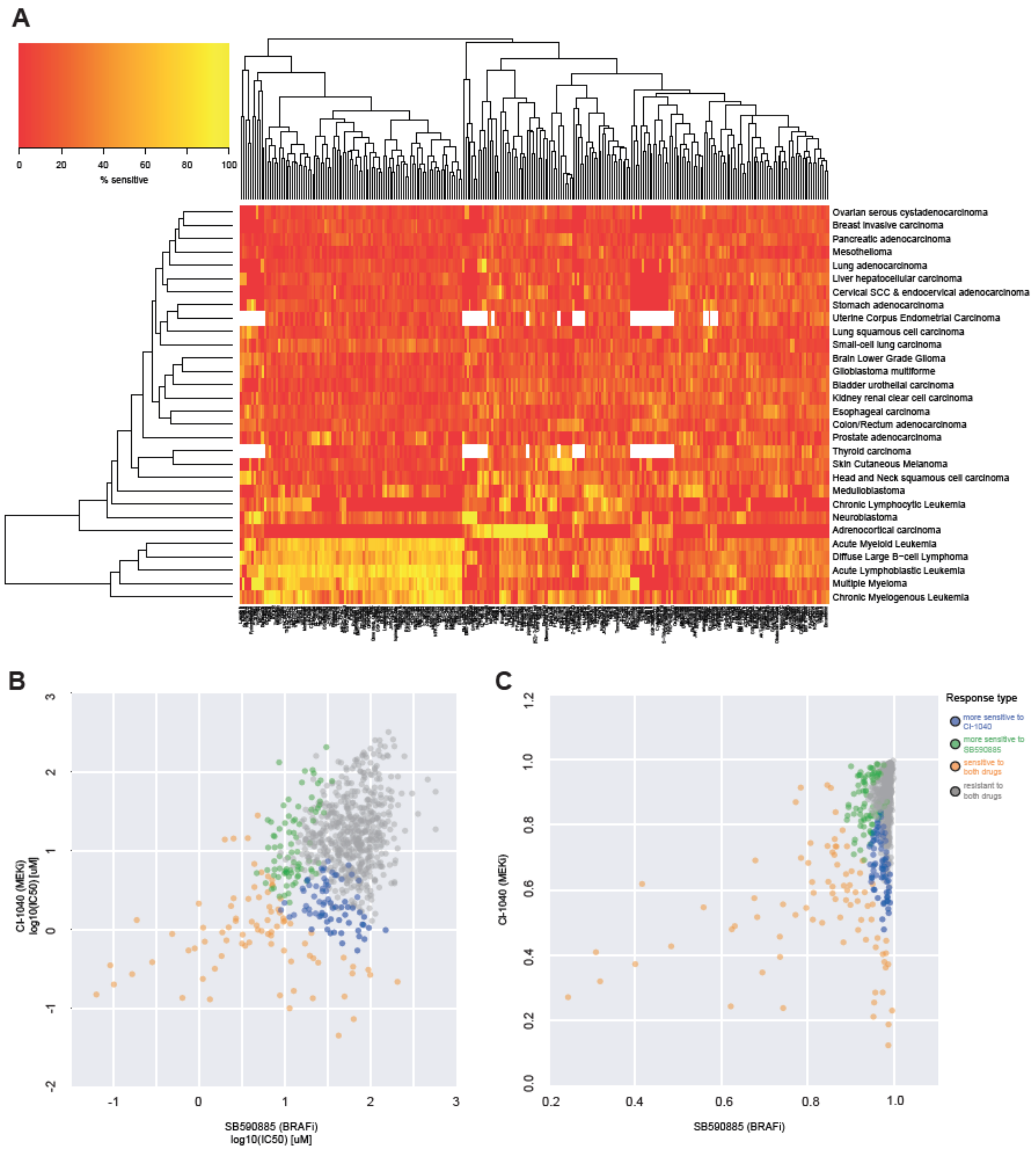


**Figure S1 Differences in evaluating drug response individually and by comparing two drugs through segmentation. (A)** Heatmap showing the global monotherapy response within 30 different cancer types (vertical axis) treated with 265 drugs (horizontal axis). The range of colours illustrates the percentage of cell lines of a particular cancer type that are sensitive to a given drug, with red being a lower percentage and yellow being a higher percentage. **(B)** Scatter plot showing the log10($IC_{50}$) values of individual cell lines coloured by the subpopulation they belong to (**Figure 1C**). Segmentation was carried out based on their response to a BRAF inhibitor (SB590885) and a MEK inhibitor (CI-1040). **(C)** Similar to **(B)** but showing the AUC values of individual cell lines.

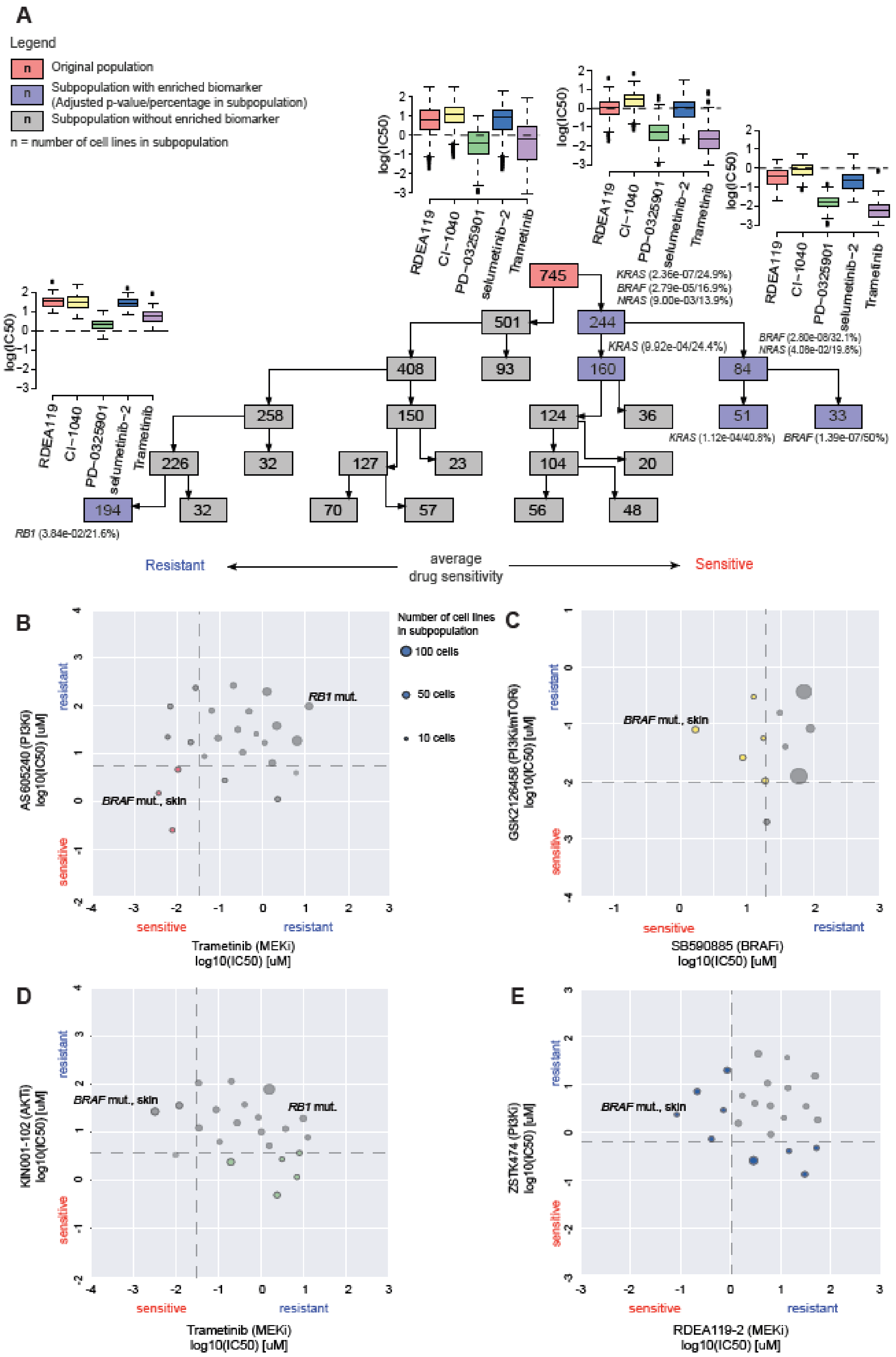
A
Legend
Original population
Subpopulation with enriched biomarker (Adjusted p-value/percentage in subpopulation)
Subpopulation without enriched biomarker
n = number of cell lines in subpopulation
RDEA119
CI−1040
PD−0325901
selumetinib−2
Trametinib
log(IC50)
745
501
244
408
93
160
84
258
150
124
36
51
33
226
32
127
23
104
20
194
32
70
57
56
48
KRAS (2.36e-07/24.9%)
BRAF (2.79e-05/16.9%)
NRAS (9.00e-03/13.9%)
KRAS (9.92e-04/24.4%)
BRAF (2.80e-08/32.1%)
NRAS (4.08e-02/19.8%)
KRAS (1.12e-04/40.8%)
BRAF (1.39e-07/50%)
RB1 (3.84e-02/21.6%)
Resistant
average drug sensitivity
Sensitive
B
Number of cell lines in subpopulation
100 cells
50 cells
10 cells
RB1 mut.
BRAF mut., skin
AS605240 (PI3Ki) log10(IC50) [uM]
resistant
sensitive
Trametinib (MEKi) log10(IC50) [uM]
C
BRAF mut., skin
GSK2126458 (PI3K/mTORi) log10(IC50) [uM]
SB590885 (BRAFi) log10(IC50) [uM]
D
BRAF mut., skin
RB1 mut.
KIN001-102 (AKTi) log10(IC50) [uM]
Trametinib (MEKi) log10(IC50) [uM]
E
BRAF mut., skin
ZSTK474 (PI3Ki) log10(IC50) [uM]
RDEA119-2 (MEKi) log10(IC50) [uM]

84

**Figure S2 Distribution of subpopulations with differential drug response after segmentation when comparing multiple drugs. (A)** Segmentation of pharmacological pattern of response for MAPK and AKT/PI3K pathway targets. There are five different MEK inhibitors (RDEA119-2, CI-1040, PD-0325901, selumetinib, and trametinib) showing the segmentation of 745 cell lines into subpopulations having distinct pharmacological patterns of response (see boxplots). Significance testing reveals enriched mutations in the subpopulations (blue boxes). **(B)** Scatter plot showing subpopulations (pink) sensitive to both trametinib (MEK inhibitor) and AS605240 (PI3K inhibitor). **(C)** Scatter plot showing subpopulations (yellow) sensitive to SB590885 (BRAF inhibitor) but resistant to GSK2126458 (PI3K inhibitor)**. (D)** Scatter plot showing subpopulations (green) sensitive to KIN001-102 (AKT inhibitor) but resistant to trametinib (MEK inhibitor). **(E)** Scatter plot showing subpopulations (blue) sensitive to RDEA119-2 (MEK inhibitor) but resistant to ZSTK474 (PI3K inhibitor) and sensitive to ZSTK474 but resistant to RDEA119-2.

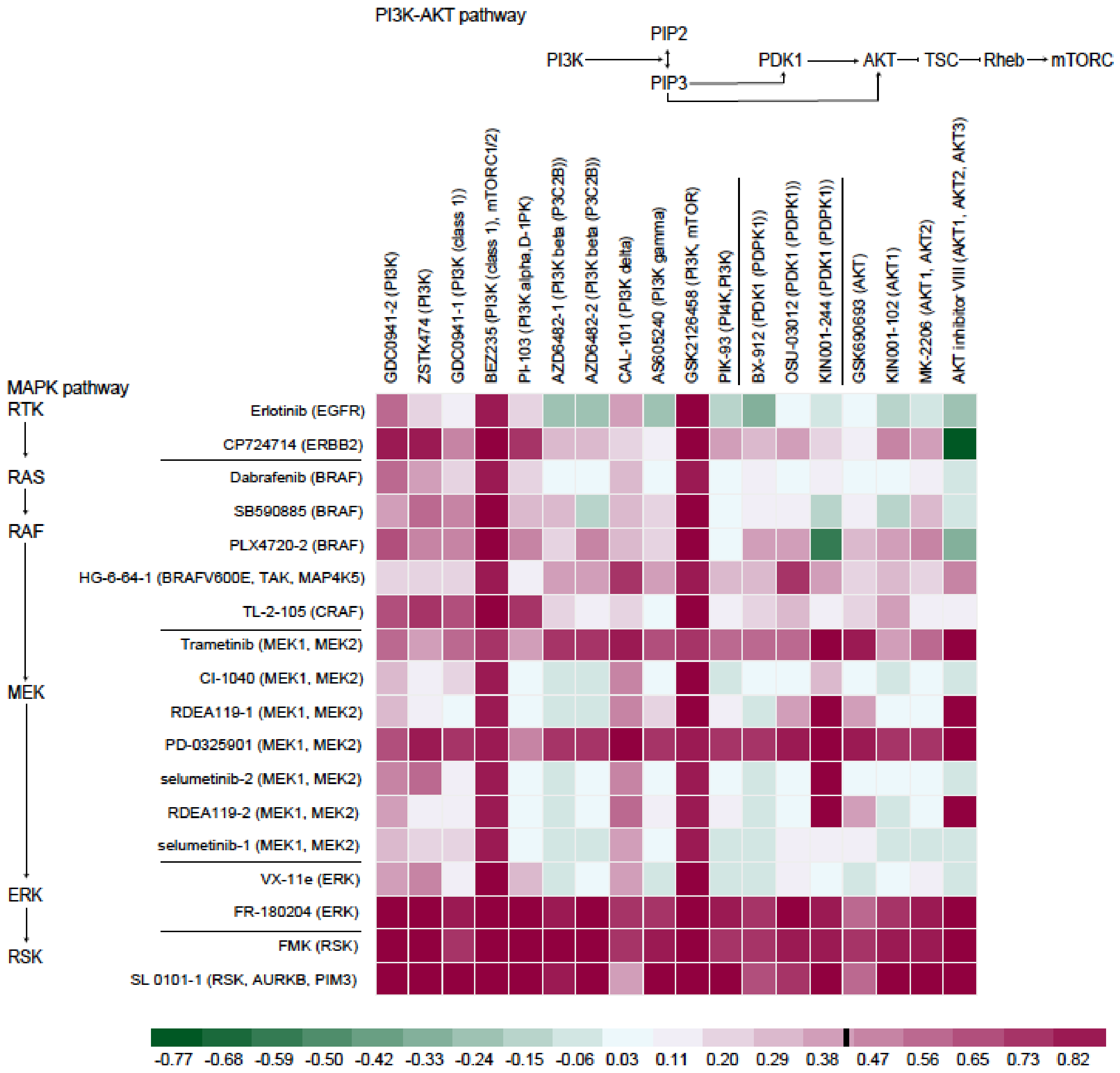


**Figure S3 Average Pearson correlation coefficient of each drug pair.** The average correlation coefficient of each drug pair was calculated by first constructing the variables vector as *x.* The average $IC_{50}$ value of the subpopulations is represented as vector *c* and the size of the subpopulations as vector *s*. We compute $z = c \circ s$ and then *c* is stacked with *z* in sequence horizontally to construct the variables vector *x*. *z* is a vector where each entry is the summation of the $IC_{50}$ value of each cell line grouped in the same subpopulation; the correlation coefficient matrix could be calculated by employing vector x. The average value in the correlation coefficient matrix is next calculated by taking the upper triangular matrix (represented as *U*) of the correlation coefficient matrix, then use a diagonal matrix which is 1 on the main diagonal to be subtracted from *U* to get the matrix *V*. Finally, the value of *V* is summed and divided by the number of unique subpopulation *l*. $l = n * (n-1)/2n$ is the number of the subpopulation resulting in the average correlation coefficient value for the drug pair.

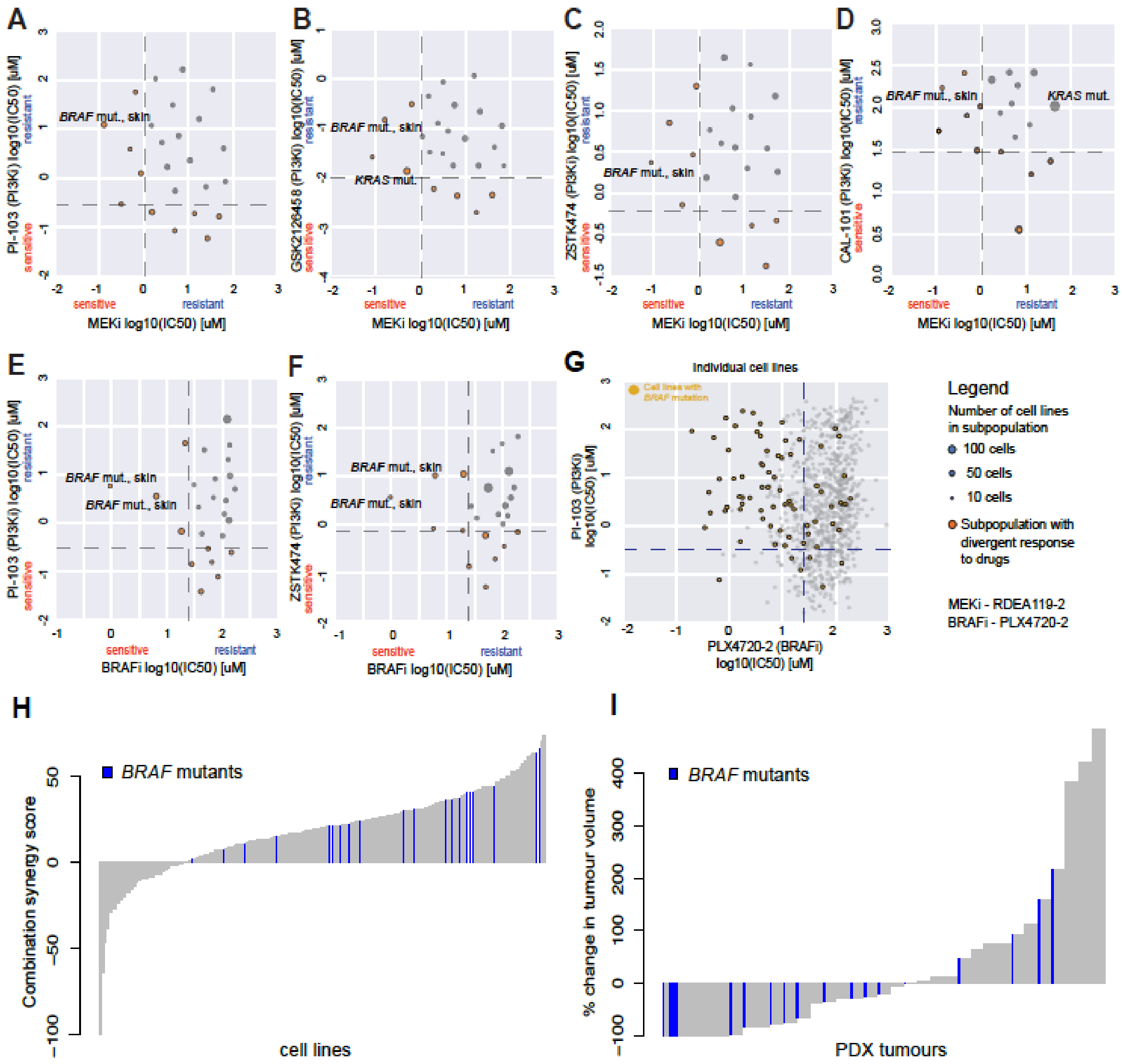


**Figure S4 Efficacy identified in drug combinations showing divergent response. (A), (B), (C), (D)** Subpopulations from comparison of response to MEK inhibitor (horizontal axis) to four different PI3K inhibitors (vertical axis) plotted by the average log($IC_{50}$) of each subpopulation. **(E), (F)** Subpopulations resulting from comparison of BRAF inhibitor (horizontal axis) to two PI3K inhibitors (vertical axis). Enriched mutations are labeled beside each subpopulation. **(G)** Scatterplot of individual cell lines without subpopulations shown in panel **(E)**. All cell lines with *BRAF* mutation are coloured gold. The dashed lines indicate the 20th percentile of log($IC_{50}$) values for each drug. **(H)** Response to combinations of MEK and

PI3K inhibitors tested in cell lines, and **(I)** BRAF and PI3K inhibitors tested in patient-derived xenograft (PDX) tumours. Higher synergy score in cell lines indicate greater response to the combination therapy, and lower % change in tumor volume indicate greater response in PDX tumors.

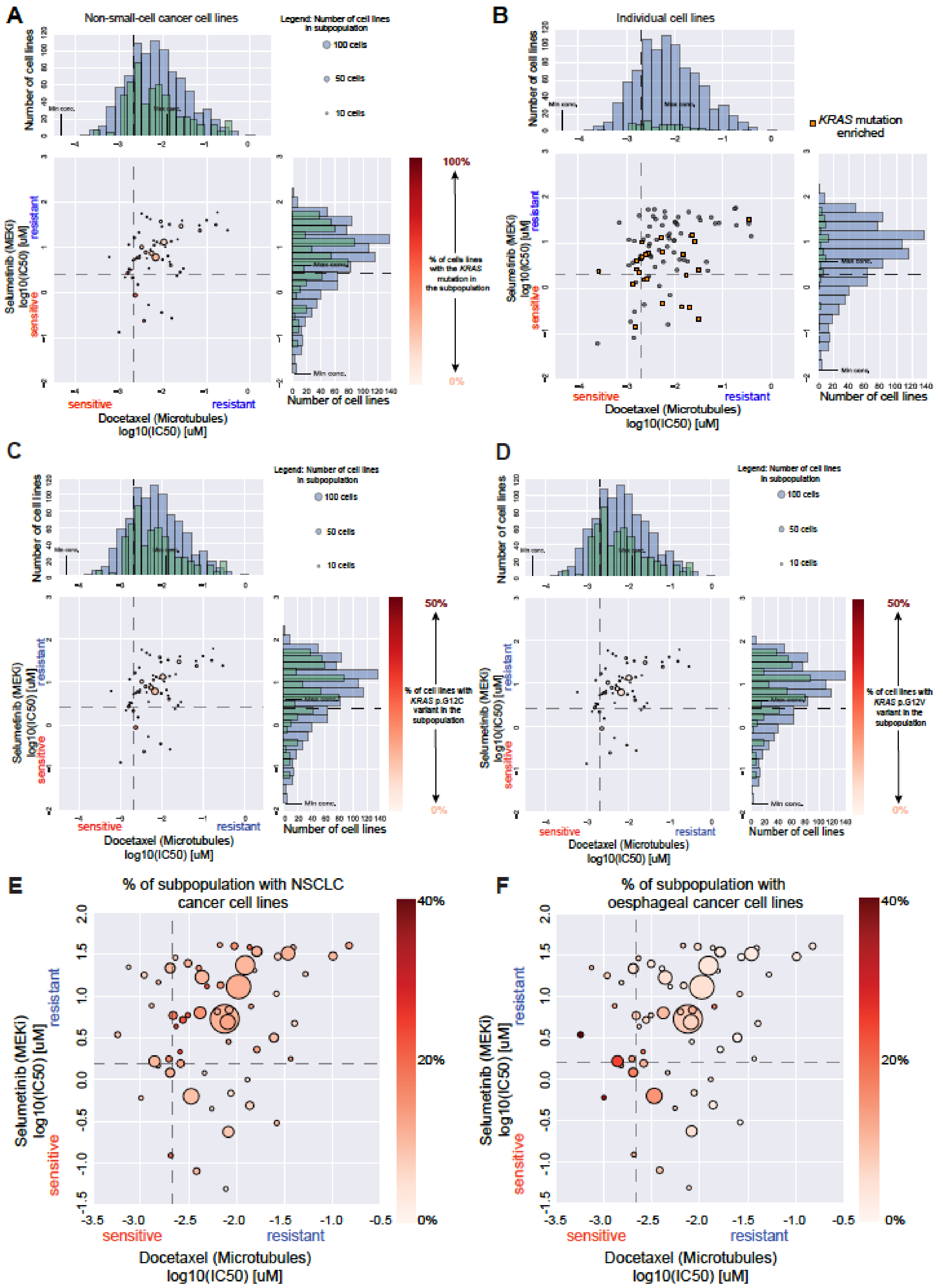


**Figure S5 Analysis of the distribution of *KRAS* mutant NSCLC cell lines treated with docetaxel (microtubules) and selumetinib (MEK inhibitor). (A)** Scatter plot showing the percentage of *KRAS* mutant NSCLC cell lines within each subpopulation. The subpopulations contain cell lines in all levels of the segmentation process. **(B)** Scatter plot illustrating the distribution of individual NSCLC cell lines treated with docetaxel and selumetinib. These cell lines are from subpopulations at the terminal level of the segmentation process. **(C)** Scatter plot showing the percentage of NSCLC cell lines with the *KRAS* p.G12C cell lines within each subpopulation. **(D)** Scatter plot showing the percentage of NSCLC cell lines with the KRAS p.G12V cell lines within each subpopulation. **(E)** Scatter plot of subpopulations illustrating the percentage of NSCLC cell lines in each subpopulation of pan-cancer cell lines. **(F)** Same as panel **(E)** but for aerodigestive cancer cell lines.

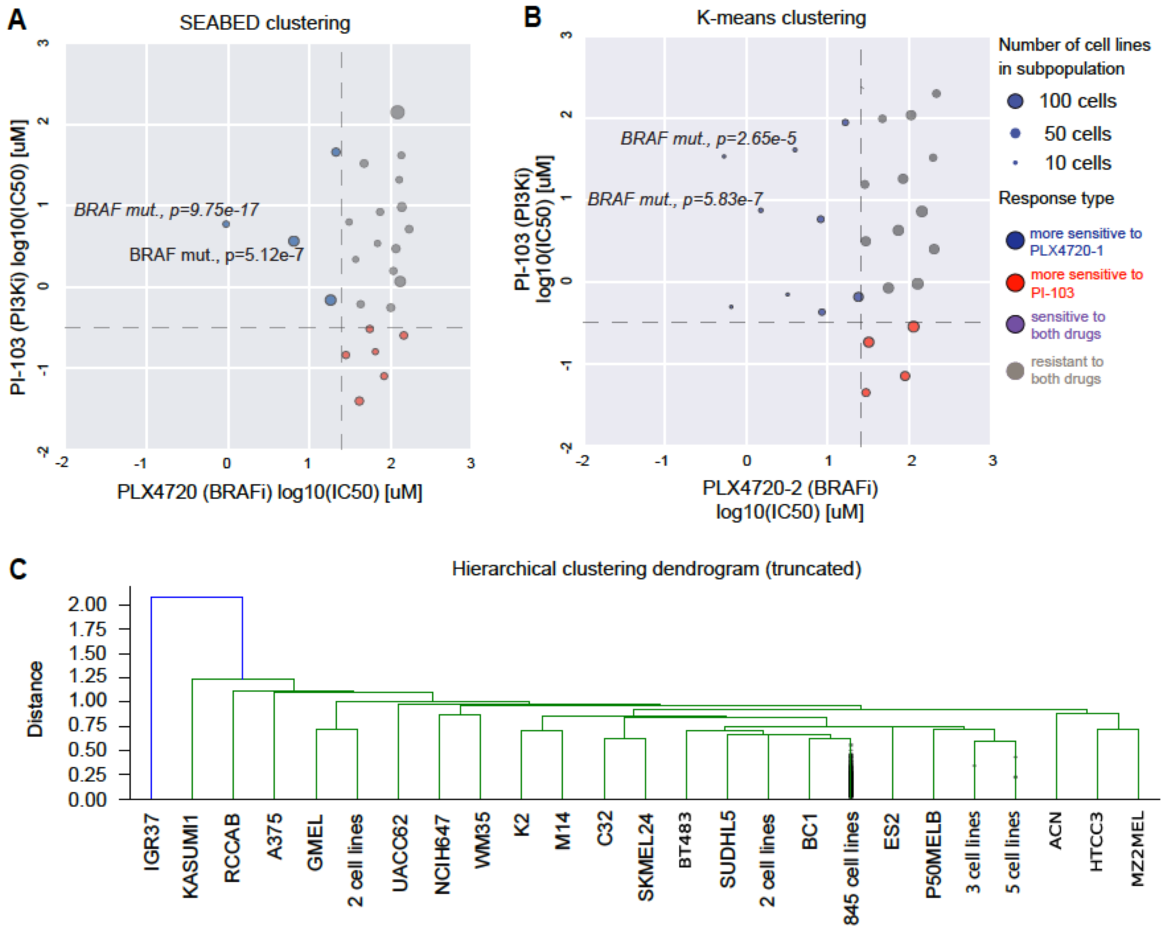


**Figure S6 Comparison of clustering approaches for a drug pair. (A)** Subpopulations and biomarkers produced by SEABED for the same drug pair (PLX4720 and PI-103). **(B)** Subpopulations identified using K-Means clustering for the same drug pair and biomarkers detected using the same technique. **(C)** Subpopulations identified using hierarchical clustering for the same drug pair. 20 of the clusters produced by hierarchical clustering contained only one cell line. Both K-Means and hierarchical were set to generate the same number of clusters as SEABED.

### Supplementary Tables

| | Selumetinib + Docetaxel | Placebo + Docetaxel | Hazard Ratio | P-value |
|---|---|---|---|---|
| Participants (KRAS mutation positive) Analyzed | 254 | 256 | | |
| Median Progression-Free Survival (Inter-Quartile Range) | 3.9 (1.5 to 5.9) | 2.8 (1.4 to 5.5) | 0.93 | 0.4355 |
| Median Overall Survival (Inter-Quartile Range) | 8.7 (3.6 to 16.8) | 7.9 (3.8 to 20.1) | 1.05 | 0.6431 |

**Table S1** Primary and secondary endpoints for the SELECT-1 Trial evaluating the efficacy of selumetinib and docetaxel in locally advanced or metastatic NSCLC (Jänne et al., 2017).

| | K-Means | Hierarchical Clustering | SEABED |
|---|---|---|---|
| **Minimum** | 4 | 1 | 20 |
| **Median** | 36 | 1 | 33 |
| **Maximum** | 81 | 846 | 83 |

**Tabel S2** Minimum, median and maximum of subpopulation sizes for different segmentation techniques for PLX4720-2 (BRAF inhibitor) and PI-103 in Figure 3.

| | K-Means | Hierarchical Clustering | SEABED |
|---|---|---|---|
| **Silhouette Coefficient** | 0.323 ± 0.037 | 0.036 ± 0.018 | 0.231 ± 0.056 |
| **RMSSTD** | 0.228 ± 0.135 | 0.854 ± 0.050 | 0.198 ± 0.103 |

**Tabel S3** Mean pairwise cluster validation results for two different for different segmentation techniques for PLX4720-2 (BRAF inhibitor) and PI-103 in Figure 3. Larger silhouette values indicate greater separation, while smaller values of RMSSTD indicate higher cluster homogeneity.

### Key Resources Table

| REAGENT or RESOURCE | SOURCE | LOCATION |
|---|---|---|
| Deposited Data | | |
| The Genomics of Drug Sensitivity in Cancer (GDSC) database | (Garnett et al., 2012; Iorio et al., 2016) | https://www.cancerrxgene.org/ |
| Cancer Cell Line Encyclopedia (CCLE) | (Barretina et al., 2012) | https://portals.broadinstitute.org/ccle |
| Cancer Therapeutics Response Portal (CTRP) | (Basu et al., 2013; Rees et al., 2016; Seashore-Ludlow et al., 2015) | https://portals.broadinstitute.org/ctrp/ |
| Menden et al. dataset | (Menden et al.) | |
| Gao et al. dataset | (Gao et al., 2015) | |
| Raw and analysed datasets | This paper | Tables S4 and S5 |
| Software and Algorithms | | |
| SEABED code | https://github.com/szen95/SEABED | N/A |
| Matplotlib | (Hunter, 2007) | N/A |
| Numpy | (Oliphant, 2006) | N/A |
| Pandas | (McKinney, 2017) | N/A |
| Bioconductor (R) | www.bioconductor.org | N/A |

| | | |
|---|---|---|
| graph-tool | https://graph-tool.skewed.de | N/A |
